# Suppression and facilitation of human neural responses

**DOI:** 10.1101/174466

**Authors:** M-P. Schallmo, A.M. Kale, R. Millin, A.V. Flevaris, Z. Brkanac, R.A.E. Edden, R.A. Bernier, S.O. Murray

## Abstract

Efficient neural processing depends on regulating responses through suppression and facilitation of neural activity. Utilizing a well-known visual motion paradigm that evokes behavioral suppression and facilitation, and combining 5 different methodologies (behavioral psychophysics, computational modeling, functional MRI, pharmacology, and magnetic resonance spectroscopy), we provide evidence that challenges commonly held assumptions about the neural processes underlying suppression and facilitation. We show that: 1) both suppression and facilitation can emerge from a single, computational principle – divisive normalization; there is no need to invoke separate neural mechanisms, 2) neural suppression and facilitation in the motion-selective area MT mirror perception, but strong suppression also occurs in earlier visual areas, and 3) suppression is not driven by GABA-mediated inhibition. Thus, while commonly used spatial suppression paradigms may provide insight into neural response magnitudes in visual areas, they cannot be used to infer neural inhibition.

## Introduction

Processes that regulate the level of activity within neural circuits^1^ are thought to play a critical role in information processing by enabling efficient coding^4^. Both suppression and facilitation of neural responses are well-known to emerge in the visual system via spatial context effects, and have a variety of perceptual consequences. For example, the perception of visual motion has been reliably shown to depend on the size and contrast of a stimulus^5, 7^. Specifically, more time is needed to discriminate the direction of motion of a large high contrast grating compared to one that is small. This seemingly paradoxical effect is referred to as spatial suppression, and has been suggested to reflect GABAergic inhibitory interactions from extra-classical receptive field (RF) surrounds (Figure 1A & B). The effect of size on duration thresholds is reversed for a low contrast stimulus – less time is needed to discriminate motion direction for a large compared to small stimulus. This facilitation of behavior is referred to as spatial summation, and has been suggested to reflect neural enhancement from RF surrounds (e.g., glutamatergic excitation) and/or an enlargement of RFs at low contrast (Figure 1C).

**Figure 1.**
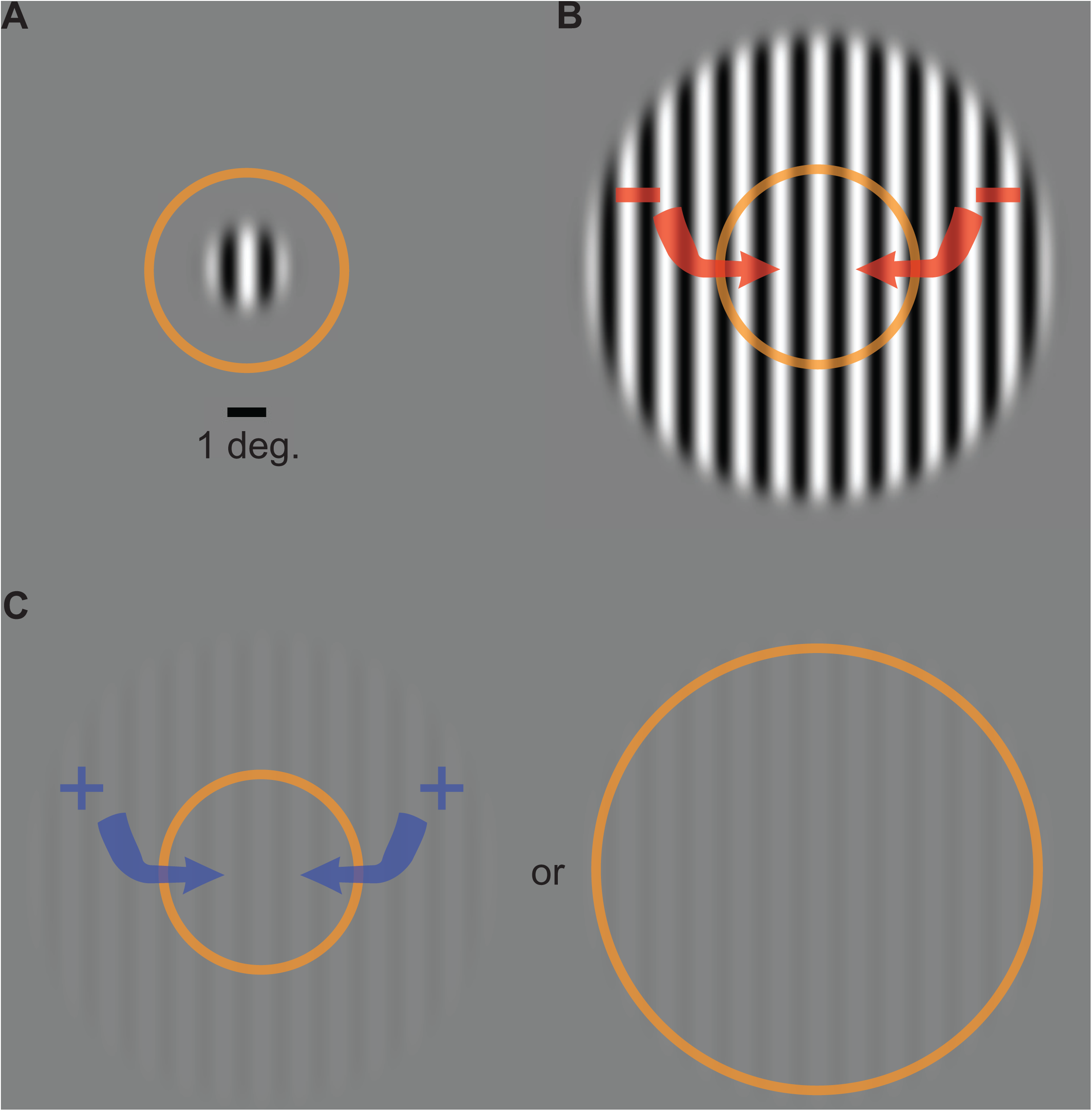
Common assumptions. The direction of motion of a small stimulus (**A**; contrast = 98%, diameter = 2°) can be perceived after a shorter presentation duration than a larger stimulus (**B**; diameter = 12°). This has been suggested to reflect the inhibitory influence of the extra-classical RF surround (red arrows) in motion-sensitive neurons in MT. Suppression turns to facilitation at low contrast (**C**; 3%), which has been assumed to reflect excitation from the surround and/or expansion of the classical RF. Orange ring represents the size of RFs in the foveal region of MT as measured in macaques^2, 3^. Comparable RF sizes in human MT are assumed^5, 6^.

Strong assumptions are often made about the neural processes underlying these seemingly complex interactions between size and contrast during motion perception. Moreover, this paradigm has been applied to the study of multiple clinical phenomena including schizophrenia^8^, major depressive disorder^9^, migraine^10^, autism spectrum disorder^11-13^, epilepsy^14^, and Alzheimer’s disease^15^, as well as normal aging^16^ and ethanol intoxication^17^. Conclusions about how neural processing is altered in these conditions have been drawn based on differences in duration thresholds relative to those of control observers. Generally, it has been assumed that spatial suppression and summation reflect distinct neural mechanisms that rely on inhibitory and excitatory processes, respectively, within brain regions involved in visual motion processing (particularly area MT).

Here we test these assumptions directly and show that: 1) spatial suppression and summation in fact naturally emerge from a single, well-established neural computation observed in visual cortex – divisive normalization^18, 19^; there is no need to posit separate mechanisms. 2) While neural responses in human MT complex (hMT+) indexed with fMRI correspond well with the measured perceptual effect, there is substantial suppression in earlier visual areas; thus, it is possible that hMT+ ‘inherits’ suppression from earlier stages of processing. 3) Two separate methodologies – magnetic resonance spectroscopy (MRS) and pharmacological potentiation of GABA_A_ receptors – demonstrate that spatial suppression is not directly linked to neural inhibition. While we find that inhibition plays a role in motion perception, increases in duration threshold as a function of stimulus size should not be taken as an index of inhibitory processing. In total, our results suggest that a single computational principle – divisive normalization – can account for spatial context effects, and that suppressive context effects are not driven by neural inhibition.

## Results

### Quantifying behavior

To quantify spatial suppression and summation psychophysically, we measured motion duration thresholds (see Methods) for 10 subjects in each of 6 different stimulus conditions, with sinusoidal luminance gratings at 3 different sizes (small [s], medium [m], and big [b]; diameter = 1, 2 & 12°, respectively) and 2 different contrasts (low = 3%; high = 98%; Figure 2A-C). The effect of stimulus size was quantified using a size index (SI; computed using the difference in thresholds between small and larger size conditions; see Methods, Equation 2). Negative SI values indicate more time was needed for motion discrimination with larger stimuli (spatial suppression), while positive values indicate shorter durations for larger stimuli (spatial summation). As expected^5, 7^, SIs depended on both size and contrast (*F*_2,9_ = 27.3, *p* = 4 x 10^-6^), with spatial suppression observed at high contrast and spatial summation at low contrast (Figure 2D).

**Figure 2.**
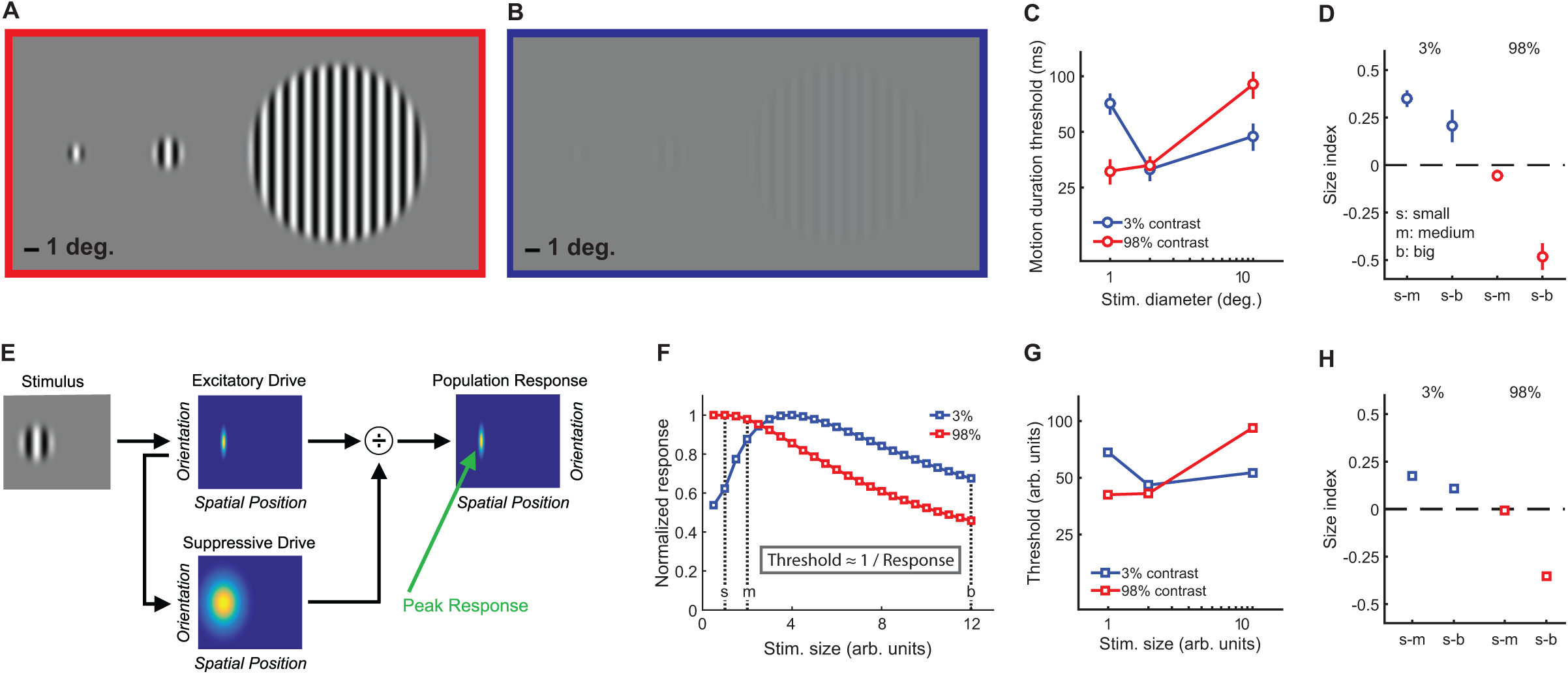
Stimuli, psychophysical results, and modeling. Small, medium, and big stimuli at high (**A**) and low contrast (**B**). The amount of time required to discriminate left- vs. right-moving stimuli with 80% accuracy (threshold in ms) is shown in **C** (average across N = 10 subjects, error bars are mean ± s.e.m.). Size indices (**D**) show the effect of increasing stimulus size, where negative values indicate that thresholds increase (suppression) and positive values indicate decreased thresholds (summation). A schematic representation of the normalization model is presented in **E** (for full model details, see Supplemental Information), with the peak predicted responses for different stimulus sizes and contrasts shown in **F** (responses for both contrasts normalized to a maximum value of 1). As noted in the inset, predicted thresholds for motion discrimination are inversely proportional to these peak responses. Thresholds (**G**) and size indices (**H**) predicted by the model show a good qualitative match to the psychophysical data (**C** & **D**).

### A single computational framework for suppression and summation

The psychophysical effects of spatial suppression and summation – sometimes attributed to distinct neural mechanisms (Figure 1) – appear to depend on a complex interplay between stimulus size and contrast. We examined whether this apparent complexity could be explained by a simple, well-established model of early visual cortical responses that incorporates divisive normalization^18, 19^, which can be summarized as:

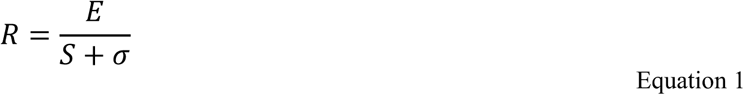

This model describes the response (*R*) to a visual stimulus in terms of an excitatory drive term (*E*; reflecting the strength of the input), divided by the sum of a suppressive drive term (*S*; also depends on input strength, but is spatially broader; Figure 2E) plus a small number known as the semi-saturation constant (*σ;* controls response sensitivity; see Methods and Supplemental Information for further model details). We found that a good qualitative match between our psychophysical data and the model predictions (Figure 2F-H; Supplemental Table 1) could be obtained by assuming that the amount of time required for motion discrimination is inversely related to the peak modeled response (see Methods; Equation 3).

At high contrast, as stimulus size grows the modeled suppressive drive increases relative to the excitatory drive, so the predicted response decreases (i.e., spatial suppression; red curve in Figure 2F). This pattern generally holds as long as the spatial tuning width of the excitatory drive is comparable with the smaller stimulus sizes, and the suppressive spatial tuning is larger than this. For low contrasts and small sizes, the suppressive drive is relatively weak compared to the semi-saturation constant (*σ*), so the response to low contrast stimuli increases with stimulus size until the suppressive drive is relatively larger than *σ* (i.e., spatial summation; blue curve in Figure 2F). Note that the parameters that determine spatial selectivity (i.e., model receptive field size) do not vary with stimulus contrast. Instead, within the computational framework of this model, spatial suppression vs. summation depends on the strength of the suppressive drive relative to *σ* at a given stimulus contrast. In general, given comparable spatial parameters to those used here, summation is predicted for values of *σ* that are within about 2 orders of magnitude of the stimulus contrast. In macaque visual cortex, the peak neural response is likewise often observed at larger sizes for lower contrast stimuli^20, 21^. Thus, the complex interplay between stimulus contrast and size that results in spatial suppression and summation during motion perception may be accounted for by a simple divisive normalization rule – it is not necessary to invoke distinct neural mechanisms to explain both phenomena.

### Neural responses reflect suppression and summation

Both our modeling work and previous studies^3, 7, 22^ suggest that spatial suppression and summation depend on reduced and enhanced neural responses (respectively) within visual cortex, particularly in area MT. To test this hypothesis directly in human visual cortex, we measured fMRI responses to low and high contrast moving gratings at different sizes with the same 10 subjects from the psychophysical experiment above. In a blocked experimental design (Figure 3A), we measured the change in the fMRI response evoked by increasing stimulus size. We report the responses from two areas: 1) early visual cortex (EVC; Supplemental Figure 1A) near the occipital pole, corresponding to the foveal confluence adjoining areas V1, V2, & V3, and 2) human MT complex (hMT+; Supplemental Figure 1C). For both EVC and hMT+, we used an independent localizer scan to define sub regions-of-interest (sub-ROIs) corresponding to cortical areas that selectively respond to the smallest stimulus size. During a single experimental scan, two stimulus sizes were presented in alternating 10 s blocks; either small and medium or small and big. Thus, the sub-ROIs – defined by the smallest stimulus size – were constantly stimulated during the scan. If the response in these sub-ROIs did not change between the small and larger stimuli (the null hypothesis), this would indicate no influence of surrounding regions. Increased responses would reflect spatial summation, while spatial suppression would yield response decreases.

**Figure 3.**
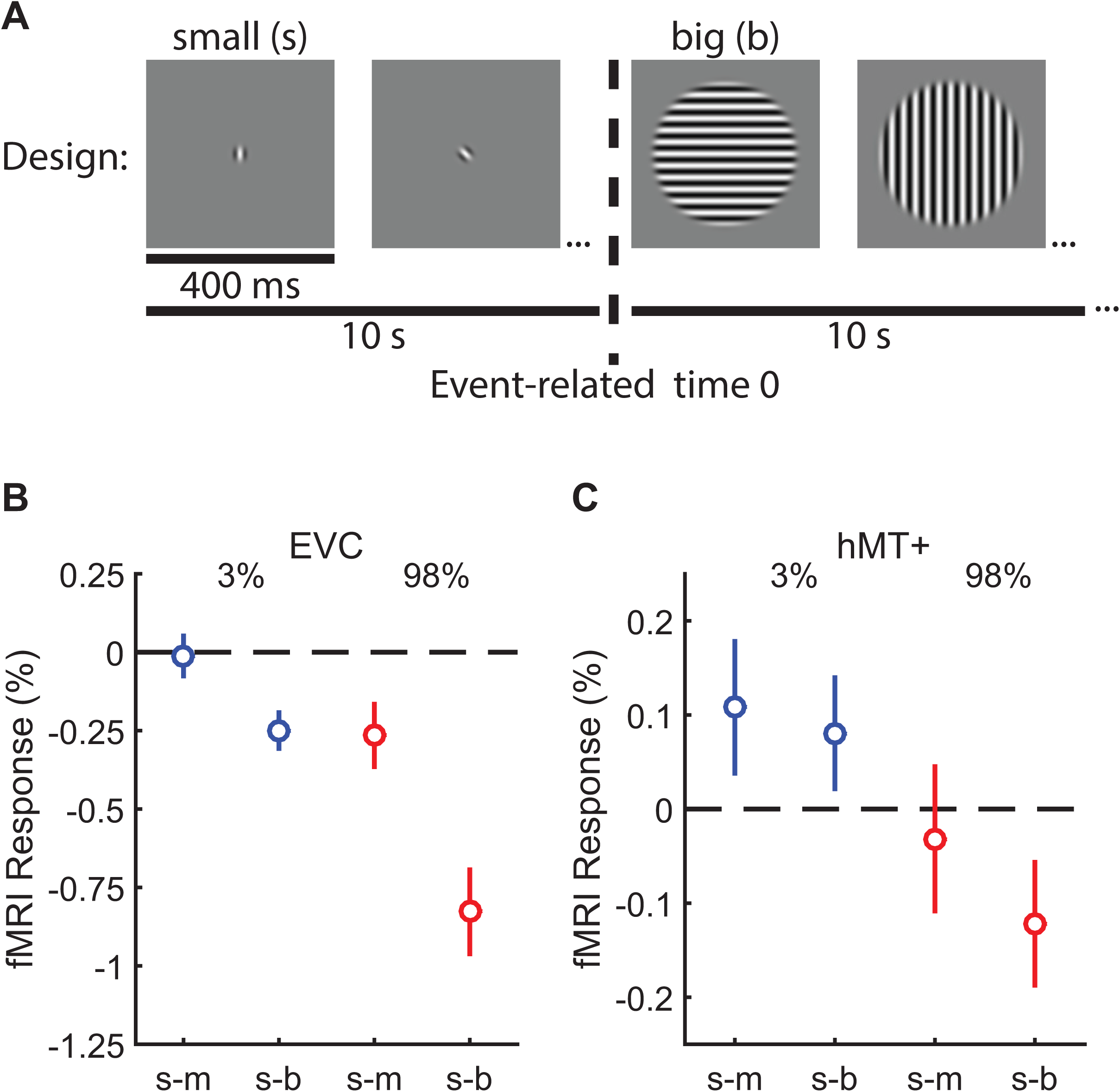
Measuring suppression and summation using functional MRI. This experiment measured the response to increasing stimulus size within regions of visual cortex representing the smallest stimulus. ROIs were localized in N = 10 subjects in EVC and N = 8 in hMT+. The blocked experimental design is illustrated in **A**. Drifting gratings (400 ms on, 225 ms blank) of a particular size were presented within 10 s blocks. In **B**, we show the change in the fMRI response within EVC and hMT+ following the increase in stimulus size from small to medium (s-m) or small to big (s-b). The response to low contrast stimuli (3%) is shown in blue, high contrast (98%) in red. Error bars are mean ± s.e.m.

In EVC there was no evidence for neural summation at low contrast; responses did not change (small-to-medium [s-m]; Supplemental Figure 1B) or decreased slightly (s-b) when low contrast stimuli became larger. However, at high contrast, suppression was observed in EVC (*F*_1,9_ = 18.0, *p* = 0.002) and was particularly strong in the small-to-big condition. In hMT+, there was evidence for both neural summation and suppression at low and high contrast, respectively. FMRI responses in hMT+ increased with stimulus size at low contrast (consistent with neural summation), and decreased at high contrast (consistent with neural suppression; Figure 3C; Supplemental Figure 1D; *F*_1,7_ = 9.0, *p* = 0.020). This pattern of fMRI responses is a better match to the spatial summation and suppression observed using psychophysics (from Figure 2D), as compared to EVC which did not show any summation. The overall smaller fMRI response modulation in hMT+ vs. EVC was expected due to larger receptive fields in hMT+^6^, which reduce the retinotopic selectivity of the hemodynamic response within this ROI. These fMRI results are consistent with the proposal that increased and decreased neural activity within hMT+ contributes to spatial summation and suppression (respectively) during motion perception^3, 7^. Observing both suppression and summation together within a single region is consistent with the framework of the normalization model (Figure 2H). In addition, it should be emphasized that suppression is clearly strong within earlier visual areas (EVC), even for low contrast stimuli.

### The relationship between spatial suppression and GABA-mediated inhibition

After finding a match between neural responses in visual cortex and behavioral performance in this paradigm, we asked whether spatial suppression might be driven directly by GABAergic inhibition. Here we used two separate methodologies: pharmacological potentiation of GABA_A_ receptors with the benzodiazepine lorazepam, and measurements of individual differences in GABA concentration with magnetic resonance spectroscopy (MRS). Our *a priori* hypothesis was that if suppression depends on GABA-mediated inhibition, increases in GABA signaling (either pharmacological or by natural variation in GABA concentration across individuals) should correspond with increased duration thresholds specifically for large stimuli – where suppression is greatest. We did not have a strong *a priori* hypothesis about how GABA signaling might affect duration thresholds for small stimuli, where suppression is minimal; however, our general intuition was that duration thresholds for small stimuli would be little affected.

#### Pharmacologically enhanced inhibition

We first examined the effect of potentiating inhibition at the GABA_A_ receptor through administration of the benzodiazepine drug lorazepam^23^. This compound causes chloride channels to open more readily in response to GABA binding, which yields greater inhibition of neural action potentials. In a double-blind, placebo-controlled crossover experiment, 15 subjects received either a 1.5 mg dose of lorazepam or placebo during separate experimental sessions on different days (order randomized and counter-balanced across subjects). Participants then completed the psychophysical paradigm used to examine spatial suppression and summation. This allowed us to test the hypothesis that strengthening inhibition would lead to increased spatial suppression during motion perception.

We found that increasing inhibition via lorazepam did not lead to stronger suppression – in fact, we observed weaker spatial suppression (SIs were less negative overall) under lorazepam vs. placebo (Figure 4C; *F*_1,14_ = 6.51, *p* = 0.023). Rather than increasing suppression, we found that lorazepam affected motion discrimination in two ways: first, lorazepam slightly increased thresholds in all conditions (*F*_1,14_ = 9.83, *p* = 0.007). More importantly, the effect of lorazepam (drug minus placebo) was stronger for smaller stimuli (Figure 4D; *F*_1,14_ = 6.91, *p* = 0.020) at both low and high contrast. By increasing thresholds for smaller stimuli, lorazepam effectively weakened spatial suppression (i.e., reduced the difference in thresholds between small and larger stimuli). Thus, we found that potentiating inhibition at the GABA_A_ receptor via lorazepam decreased, rather than increased spatial suppression in this paradigm. This result is not consistent with the idea that spatial suppression is directly mediated by neural inhibition.

**Figure 4.**
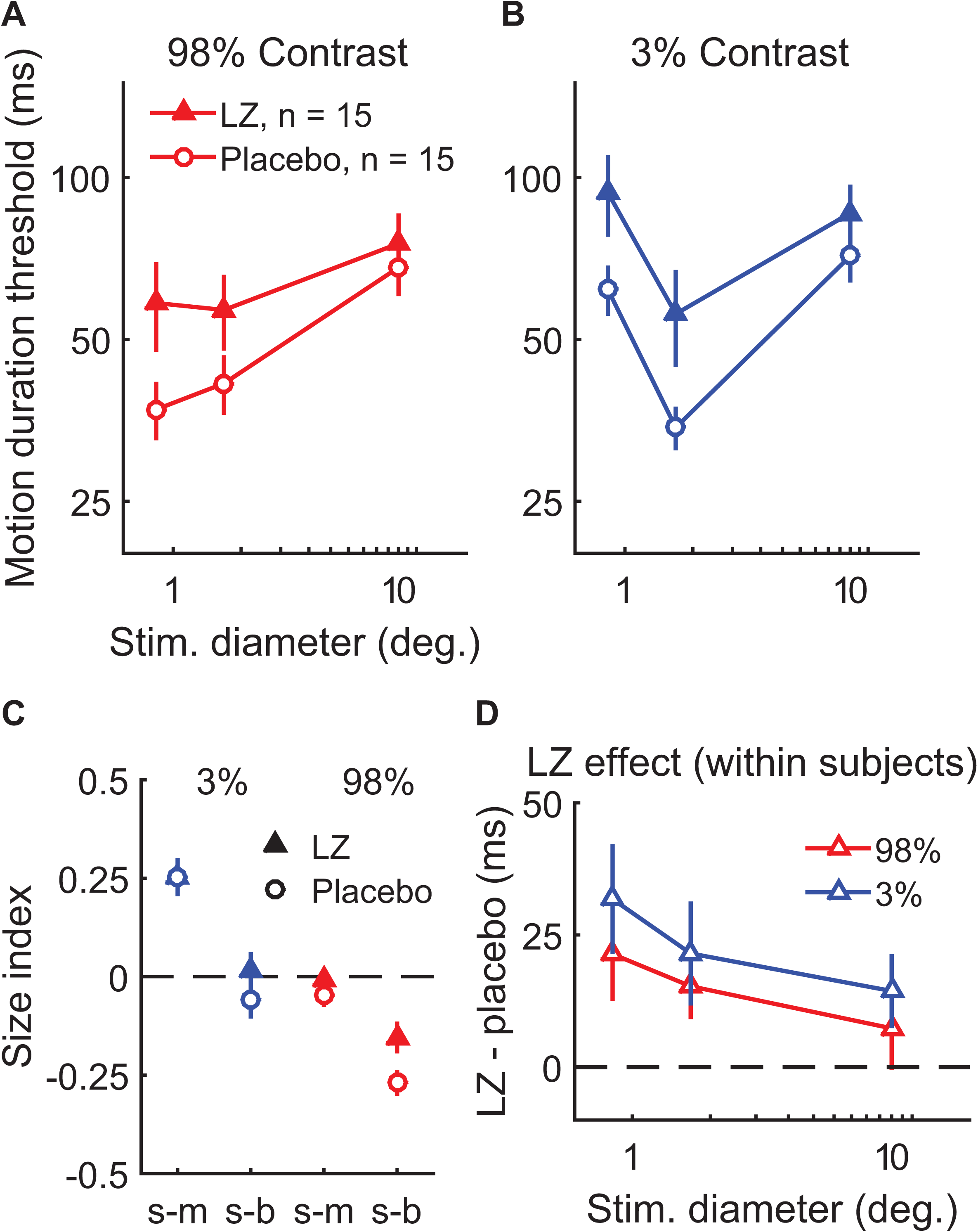
The effect of lorazepam on spatial suppression. Fifteen subjects took lorazepam (LZ) or placebo in a double-blind, cross-over experiment. Duration thresholds were measured for high (**A**) and low contrast (**B**) moving gratings in each session. Size indices (**C**) and within-subjects effects of the drug (**D**) were calculated. Error bars are mean ± s.e.m.

#### Measuring GABA via spectroscopy

To further examine the relationship between spatial suppression and inhibition, we measured the concentration of GABA+ (GABA plus co-edited macromolecules) within specific regions of visual cortex using MRS. GABA+ was quantified within moderately-sized voxels (27 cm^3^) centered on functionally-identified hMT+ (left & right hemispheres measured in separate scans and averaged; Supplemental Figure 2A), anatomically-identified EVC (average of 2 measurements in the same region; Supplemental Figure 2B), and a control voxel in anatomically-identified parietal cortex (Supplemental Figure 2C). MRS data were acquired at rest (i.e., subjects were not asked to perform a specific task). The same subjects also participated in separate psychophysical and fMRI experiments to measure spatial suppression, comparable to those described above (N = 22 complete data sets, see Supplemental Figure 3 for a summary of these results). Unlike manipulating GABA pharmacologically, MRS measurements of GABA+ are thought to reflect the stable, individual differences in baseline concentration of this neurotransmitter^24^. Measuring this trait using MRS allowed us to test the hypothesis that subjects with more GABA+ in visual cortex would show greater spatial suppression.

Behaviorally, the strength of spatial suppression did not depend on the concentration of GABA+ in hMT+; no correlations were found between hMT+ GABA+ and SIs (Supplemental Figure 4A-F; all |*r*_20-25_| < 0.32, uncorrected *p*-values > 0.14). Further, there was no association between fMRI measurements of spatial suppression in hMT+ and the concentration of GABA+ in this region (Supplemental Figure 4G-H; |*r*_19_| < 0.29, uncorrected *p*-values > 0.20). To aid visualization, in Figure 5A-C we show the psychophysical data with subjects split into two groups based on the concentration of GABA+ within hMT+ (median split). Surprisingly, we observed that more GABA+ in hMT+ predicted overall better psychophysical performance (lower thresholds on average) during motion direction discrimination (Figure 5D; *r*_20_ = -0.46, *p* = 0.030, uncorrected). These results indicate that more GABA in hMT+ is associated with better motion perception in general, but not with the strength of spatial suppression.

**Figure 5.**
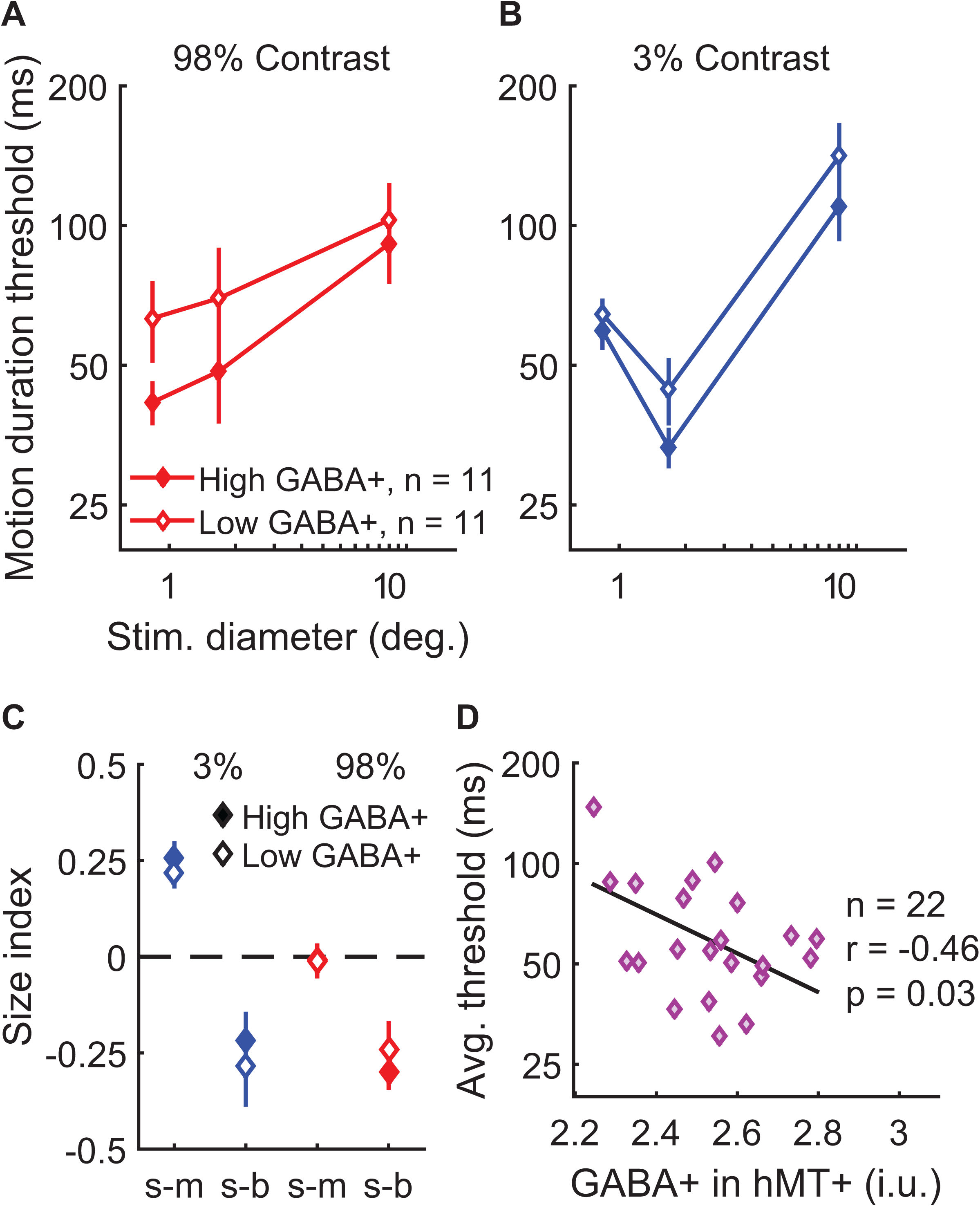
Examining task performance in terms of individual differences in GABA+ concentration in hMT+. To help illustrate the relationship between GABA+ measured in hMT+ and motion discrimination performance, thresholds (**A** & **B**) and size indices (**C**) are shown for subjects with lower (open symbols, N = 11) and higher GABA+ (filled symbols, N = 11; groups defined by median split). Error bars are mean ± s.e.m. As shown in **D**, subjects with more GABA+ in hMT+ performed better overall during motion discrimination (lower average thresholds; geometric mean of all 6 stimulus conditions).

When examining GABA+ measured within other areas (EVC or parietal cortex), no significant relationships were found with spatial suppression or overall psychophysical thresholds, nor did GABA+ correlate with spatial suppression measured in EVC using fMRI (all |*r*_19-25_| < 0.28, uncorrected *p*-values > 0.21; Supplemental Figure 5 & Supplemental Figure 6). These findings suggest that the relationship between motion discrimination performance and GABA is specific to hMT+, and not a more general (e.g., brain-wide) phenomenon. Together, our specroscopy results demonstrate that the concentration of GABA in visual cortex indexed by MRS does not predict the strength of spatial suppression during motion perception. Instead, higher GABA+ in hMT+ was associated with better motion discrimination performance overall. While this result differs from the effect of lorazepam on motion discrimination described above, we note that MRS reflects individual differences in GABA+ within visual brain areas that are (presumably) at homeostasis, whereas the effects of lorazepam can be attributed to a transient pharmacological strengthening of inhibition specific to the GABA_A_ receptor class.

## Discussion

This study examined a number of common assumptions about the neural processes underlying spatial suppression and summation during motion perception. Although this paradigm has been posited as a behavioral index for inhibition, we did not find a major role for GABA in determining the strength of spatial suppression. While a few studies have probed the neurochemical underpinnings of perceptual suppression in humans^17, 25-27^, much of our current insight into the neural mechanisms of suppression and inhibition has come from work in animal models. Some reports suggest that GABAergic inhibition plays a direct role in surround suppression within visual cortex^28-31^, while others have indicated that this suppression occurs via withdrawal of excitation^32-35^, that may be balanced by reduced inhibition. Our results demonstrate that in humans, spatial suppression is not directly mediated by inhibition; lorazepam weakened suppression rather than strengthening it, and MRS measurements of GABA+ in visual cortex did not predict suppression strength. Instead, our findings are more consistent with the withdrawal of excitation as a mechanism for spatial suppression. This agrees with an earlier observation that blocking GABA in macaque MT had no effect on spatial suppression^36^ (see also^37^). However, our MRS findings appear to conflict with reports that greater mid-occipital GABA was associated with stronger surround suppression during contrast perception^25, 27^. This discrepancy may be related to the methods for quantifying suppression, and / or differences in the role of inhibition during perception of contrast and motion.

While not the direct mechanism for suppression, we observed clear links between GABAergic inhibition and motion perception. We considered how these effects of GABA might be described in terms of the normalization model (in a manner other than directly mediating suppression). We found that potentiating inhibition at the GABA_A_ receptor via lorazepam decreased spatial suppression, rather than strengthening it. Supplemental Figure 7A-C shows how a reduction in both input (i.e., contrast) and output (i.e. response) gain yields predicted thresholds that mirror the observed effects of lorazepam (compare with Figure 4A-C; also see Supplemental Table 1). Specifically, higher thresholds for smaller stimuli following lorazepam may be accounted for in the normalization model by reducing the strength of the input (i.e., reduced contrast gain). Lowering contrast gain raised the predicted thresholds for small stimuli, but had little effect when stimuli were large. We therefore speculate that the effect of lorazepam in reducing spatial suppression may be consistent with reduced contrast gain within brain regions relevant to motion perception (e.g., MT).

From previous work that manipulated GABA_A_ receptor function in animal models^37, 38^, it may be expected that potentiating inhibition at this receptor would reduce neural responsiveness overall (i.e., lower response gain). Beyond increasing threshold for small stimuli, lorazepam also raised motion discrimination thresholds slightly in all conditions, an effect that is consistent with overall lower neural responses. While reducing *input* gain in the model (as above) slightly reduces the predicted thresholds in all conditions, such an effect may also be modeled by reducing *response* gain (scaling down predicted responses; Supplemental Figure 7A-C; Supplemental Table 1). This is also consistent with earlier behavioral studies showing that benzodiazepine administration generally reduces visual performance^26, 39^. However, we note that the observed effects of lorazepam were specific to threshold-level motion discrimination (especially for smaller stimuli); catch trial performance was not affected (see Methods), which argues against more general pharmacological effects (e.g., fatigue). In summary, while the effect of lorazepam during motion discrimination may be consistent with weaker contrast gain (and perhaps also lower response gain), our findings do not support the idea that spatial suppression is directly mediated by inhibition.

We additionally showed that higher baseline GABA (as measured by MRS) is associated with better, rather than suppressed motion perception. Because this correlation was unexpected, and no statistical correction for multiple comparisons was performed, this result should be interpreted with caution. To understand why more GABA in hMT+ might lead to better motion discrimination, we considered two possibilities. First, better performance (lower thresholds) might result if GABA *increased* neural activity in this region – this proposal does not seem parsimonious, given the well-established inhibitory function of GABA in the adult nervous system^40^. Alternatively, if more baseline GABA lowers the behavioral response criterion (without necessarily changing the neural response), possibly by improving neural signal-to-noise^41^, then we would expect better performance with higher GABA in hMT+. Indeed, within our model (Equation 3), reducing the response criterion has the same effect on the predicted threshold as an increased neural response. In Supplemental Figure 7D-F, we show how the effect of changing criteria can be described in terms of the normalization model (compare with Figure 4A-C; see also Supplemental Table 1). Note that changing the criterion has no effect on the strength of spatial suppression predicted by the model (Supplemental Figure 7F). This is consistent with our finding that the concentration of GABA+ in hMT+ does not influence spatial suppression measured psychophysically (Figure 5C; Supplemental Figure 4A-F) or with fMRI (Supplemental Figure 4G-H). Thus, lower response criteria with higher baseline GABA may plausibly account for our MRS data. However, as with our experiment using lorazepam, these MRS data contradict the notion that GABAergic inhibition directly determines the strength of spatial suppression. Altogether, our findings suggest that the assumption of a direct link between spatial suppression and inhibition is invalid.

We considered whether spatial suppression and summation might be described in terms of a single computation – divisive normalization – a model for early visual processing in which a neuron’s response is suppressed (divided) by the summed response of its neighbors^18, 19^. We are not the first to suggest that the normalization model may account for these phenomena; Rosenberg and colleagues^12^ used weaker normalization to explain superior motion discrimination performance in autism spectrum disorder^11^. This study provides the first direct application of this model to this paradigm in typically developing individuals under a variety of experimental conditions. In addition, earlier work has modeled spatial suppression and summation in terms of a variety of divisive^42^ or subtractive^43^ computations. Generally, these earlier models have treated “excitatory” and “suppressive” mechanisms separately (e.g., different contrast sensitivity). The normalization model^19^ used here provides a simpler account, wherein the suppressive drive is essentially the same as the excitatory drive, only broader in space and orientation (see Supplemental Information). We found that this normalization model is sufficient to explain the complex interplay between stimulus size and contrast that produces psychophysical (or neural) suppression and summation. This framework appears more parsimonious than positing separate mechanisms (neural or computational) operating at different stimulus contrasts to produce suppression and summation.

Our modeling work shows that normalization can describe spatial suppression and summation under a variety of different experimental conditions. We found good agreement between the predicted model response (Figure 2G & H), psychophysics (Figure 2C & D), and the fMRI response in hMT+ (Figure 3C). That both suppression and summation were observed in the fMRI response from a sub-region of hMT+ representing the stimulus center indicates that these two effects are co-localized in cortex, and both could plausibly occur within a single neural population^21^, which is consistent with the proposed model framework. Further, the way in which spatial suppression was affected by lorazepam (Supplemental Figure 7A-C), but not baseline GABA in hMT+ (Supplemental Figure 7D-F), can be explained in terms of the normalization model (Supplemental Table 1). It is worth noting that the lorazepam data were described by an exponential reduction of contrast gain, which we speculate may reflect the compounded reduction of neural responses across multiple stages of visual processing (e.g., retina, lateral geniculate nucleus, visual cortex) due to systemic potentiation of inhibition at the GABA_A_ receptor. Although recent work has suggested some limitations for describing spatial suppression in terms of normalization^44^, we find that this model framework is sufficiently general to describe the suppression effect across our different experiments.

Finally, we examined the extent to which suppression and summation may be attributed to modulation of neural activity within area MT. Our results generally supported this hypothesis; fMRI responses in hMT+ showed better agreement with spatial suppression and summation measured psychophysically, as compared with the responses in EVC (Figure 3B & C). Further, better motion discrimination across individuals was predicted by higher GABA+ in hMT+ (Figure 5D), but not in EVC (Supplemental Figure 5D). The idea that spatial suppression during perception is driven (at least in part) by surround suppression within MT is consistent with recent work in primate models^3^. For many neurons in primate MT, the presence of stimuli within the extra-classical receptive field surround suppresses (or enhances) the response to stimuli within the receptive field center^3, 21, 45, 46^. The proposed link between perceptual suppression and neural suppression in MT has received further (if limited) support from studies in humans^22, 47^. Using fMRI, MRS, and modeling, our findings extend this link by showing a correspondence between neural processing in hMT+ and motion discrimination performance in human subjects.

While neural processing in hMT+ seemed more closely linked to perception, strong suppression of fMRI responses was observed in EVC at both low and high stimulus contrast (Figure 3B). Feed-forward and feedback connections between EVC and MT are thought to play an important role in spatial context processing^20^; our data alone are not sufficient to determine the extent to which MT inherits suppression from EVC and / or drives this suppression via feedback. However, surround suppression in early visual areas (e.g., V1, V2) is well established in animal models^20^, and suppressed fMRI responses in these areas generally correspond with perceptual suppression during contrast judgments^48-50^. This raises the possibility that some amount of spatial suppression during motion discrimination may be attributed to neural suppression in EVC.

## Methods

### Participants

A total of 48 adult human subjects were recruited across all experiments. First, 10 subjects participated in both the initial experiments characterizing spatial suppression psychophysically and with fMRI (8 males and 2 females, mean age 30 years, *SD* = 6.4 years). Second, 15 subjects (7 males and 8 females, mean age 27 years, *SD* = 4.4 years) completed the lorazepam experiment. Four subjects participated in both of these first two sets of experiments. Third, 27 subjects (12 males and 15 females, mean age 24 years, *SD* = 3.6 years) completed the MRS experiment. Data from 15 subjects in the lorazepam experiment and 20 subjects in the MRS experiment were also included as part of another study (in preparation) that characterizes the effects of GABA on human perception.

All subjects reported normal or corrected-to-normal vision and no neurological impairments. Before enrollment in the lorazepam experiment, subjects were screened for potential drug interactions. Subjects participating in MRS reported no psychotropic medication use, no more than 1 cigarette per day within the past 3 months, no illicit drug use within the past month, and no alcohol use within 3 days prior to scanning. Subjects provided written informed consent prior to participation and were compensated for their time. All experimental procedures were approved by the University of Washington Institutional Review Board, and conformed to the ethical principles for research on human subjects from the Declaration of Helsinki.

### Visual display and stimuli

Psychophysical experiments were performed using one of two display apparatuses in different physical locations for logistical reasons: 1) a ViewSonic G90fB CRT monitor (refresh rate = 85 Hz; used for the data shown in Figure 2), or 2) a ViewSonic PF790 CRT monitor (120 Hz; used for all other experiments) with an associated Bits# stimulus processor (Cambridge Research Systems, Kent, UK). In both cases, stimuli were presented on Windows PCs in MATLAB (MathWorks, Natick, MA) using Psychtoolbox-3^51-53^, and a chin rest was used to stabilize head position. During fMRI, stimuli were displayed via projector; either an Epson Powerlite 7250 or an Eiki LCXL100A (following a hardware failure), both operating at 60 Hz. Images were presented on a semicircular screen at the back of the scanner bore, and viewed through a mirror mounted on the head coil. Stimuli during fMRI were displayed using Presentation software (Neurobehavioral Systems, Berkeley, CA). The luminance of all displays was linearized using custom software. Viewing distance was 52 cm for psychophysical display #1, and 66 cm for both display #2 and in the scanner.

In each experiment, we presented drifting sinusoidal luminance modulated gratings at two different Michelson contrast levels (low = 3%, high = 98%) and 3 different sizes (small, medium & big; see Figure 2A & B), following the method of Foss-Feig and colleagues^11^. Stimulus diameter was 1, 2 & 12° visual angle for the small, medium, and big stimuli (respectively) in all fMRI experiments and the first psychophysical experiment (data shown in Figure 2; using display #1). Due to a coding error, the stimulus diameter was slightly smaller in all subsequent psychophysical experiments (performed using display #2; diameter = 0.84, 1.7 and 10°). Drift rate was always 4 cycles/s. Gratings were presented within a circular window, whose edges were blurred with a Gaussian envelope (*SD* = 0.25° for display #1 & fMRI, 0.21° for display #2). Stimuli were presented centrally on a mean luminance background, and had a spatial frequency of 1 cycle/° (display #1 & fMRI) or 1.2 cycles/° (display #2).

### Paradigm and data analysis

#### Psychophysics

Subjects were asked to determine whether a briefly presented vertical grating drifted left or right (randomized and counterbalanced). Trials began with a central fixation mark; either a small shrinking circle (850 ms, for the MRS experiments) or a static square (400 ms, all other experiments). This was followed by a blank screen (150 ms), after which the grating stimuli appeared (variable duration, range 6.7 – 333 ms), followed by another blank screen (150 ms), and finally a fixation mark (the response cue). Subjects indicated their response (left or right) using the arrow keys. Response time was not limited. To permit very brief stimulus presentations, gratings appeared within a trapezoidal temporal envelope, following an established method^11^. Thus, the first and last frames were presented at sub-maximal contrast, and the duration was defined by the full-width at half-maximum contrast.

Duration of the grating stimuli varied across trials according to a Psi adaptive staircase procedure^54^ controlled using the Palamedes toolbox^55^. Duration was adjusted across trials based on task performance, to determine the amount of time needed to correctly discriminate motion direction with 80% accuracy (i.e., the threshold duration). Staircases were run separately to determine thresholds for each of the six stimulus conditions (2 contrasts x 3 sizes, as above). Condition order was randomized across trials. Thirty trials were run per staircase within a single run (approximately 6 min). There were also 10 catch trials per run (all large, high contrast gratings, 333 ms duration), which were used to assess off-task performance. Each subject completed 4 runs, with a total experiment duration of about 30 min. Example and practice trials were presented before the first run. For 5 subjects in the MRS experiment, thresholds were not obtained for the smallest stimulus size.

Psychometric thresholds and slopes were quantified for each run by fitting the discrimination accuracy data with a Weibull function using maximum likelihood estimation^54^. Guess and lapse rate were fixed at 50% & 4%, respectively. Threshold duration was defined at 80% accuracy based on this fit. Threshold estimates below 0 ms or above 500 ms were excluded; a total of 4 such threshold estimates were excluded across all experiments. When averaging across thresholds from different stimulus conditions (e.g., Figure 5D) we computed a geometric mean, to account for the fact that the threshold range varied across conditions. The effect of stimulus size on task performance was quantified using size indices (SIs), such that:

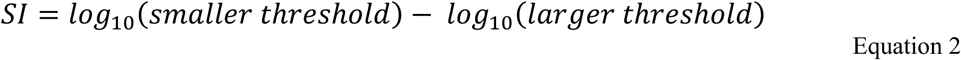

#### Computational modeling

We examined the extent to which spatial suppression could be qualitatively explained in terms of a well-established divisive normalization model^18, 19^. An equation that summarizes the model (Equation 1) is included in the Results, and a graphical depiction of the model is presented in Figure 2E. Full details of the model are provided in the Supplemental Information. Briefly, the values of *E* and *S* in Equation 1 depend on the properties of the stimulus (e.g., size, orientation, contrast), as well as a number of other parameters that determine how sensitive the model is to different stimulus properties (e.g., the spatial extent of excitation and suppression; Supplemental Equation 1 & Supplemental Equation 2). The value of *σ* determines the sensitivity of the response to weak stimuli, and prevents the response from being undefined when a stimulus is absent. These parameters are derived from an extensive literature describing how neurons in early visual cortex respond to different properties of visual stimuli^19^, and are listed for each instantiation of the model in Supplemental Table 1.

A good qualitative match between the model and our psychophysical data was obtained with minimal adjustments of the model assumptions (i.e., free parameters were adjusted manually, rather than estimated based on a computational fit to our data). We used a “winner-take-all” decision rule in reading out the population response from the model, which had the consequence of choosing the response from the center of the population being modeled (i.e., the response to the center of the stimulus, which was always the largest within the population). To equate the response predicted by the model to motion discrimination thresholds, we assumed an inverse relationship between predicted response and discrimination time, such that:

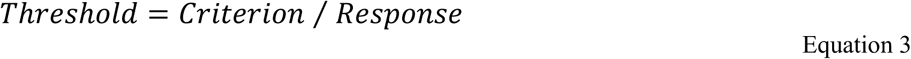

where *Threshold* is the amount of time needed to discriminate the direction of stimulus motion, *Criterion* represents an arbitrary response value that must be reached to make the perceptual judgment, and *Response* is the predicted response rate (from Equation 1). This framework is consistent both with previous modeling efforts^42, 43^, and electrophysiological work showing a close correspondence between psychophysical discrimination thresholds and neural response magnitudes in macaque MT during motion perception^3, 56^. The *Criterion* value varied across different versions of the model (Supplemental Table 1) and different contrast levels – typically, a lower *Criterion* was used to model thresholds for 3% versus 98% contrast stimuli. However, the value of *Criterion* was held constant across stimulus sizes for a given contrast.

We emphasize that our goal was not to quantitatively describe the data in terms of the model but instead to show that normalization, as a computational principle, is sufficient to qualitatively account for our findings. In general, we used parameter values that were similar to previous instantiations^19, 57^, and/or approximate the realistic values of neurons in visual cortex. Rather than make any claims about the specifics of the parameter values, we instead note in the Results the relationships between parameters (e.g., suppressive drive having broader spatial tuning than excitatory drive) that are necessary to predict the general pattern of results from our experiments.

#### Lorazepam

In separate experimental sessions separated by at least 1 week, subjects received either 1.5 mg lorazepam or placebo, with the order randomized and counter-balanced across subjects. The compounds were dispensed by a pharmacist who was not involved the study; both subjects and experimenters were blind to the order of drug & placebo until after both experimental sessions were complete. Following a 2-hour wash-in period, subjects completed the above psychophysical paradigm as part of a larger battery of experiments lasting approximately 1.5 hours. The order in which the spatial suppression paradigm was performed within this series was randomized and counter-balanced across subjects, but was always the same for the drug and placebo sessions within each subject. Instructions and practice trials were presented before the experiment in both sessions.

Catch trial accuracy was used to assess whether lorazepam affected cognitive performance in general, or motion perception more specifically. Accuracy was equivalently high in both placebo (mean = 99%, *SD* = 1.8%) and drug sessions (mean = 98%, *SD* = 3.6%; paired *t*-test, *t*_14_ = 0.8, *p* = 0.4), indicating that lorazepam specifically reduced threshold-level motion discrimination, and not task performance more generally (e.g., reduced performance due to fatigue).

#### Functional MRI

Data were acquired on a Philips Achieva 3 Tesla scanner. A T_1_-weighted structural MRI scan was acquired during each session with 1 mm isotropic resolution. Gradient echo fMRI data were acquired with 3 mm isotropic resolution in 30 oblique-axial slices separated by a 0.5 mm gap (2 s TR, 25 ms TE, 79° flip angle, A-P phase-encode direction). A single opposite direction (P-A) phase-encode scan was acquired for distortion compensation. Each scanning session lasted approximately 1 hour.

Our fMRI paradigm examined the change in the fMRI signal in response to an increase in stimulus size (e.g., spatial suppression). This involved presenting smaller and larger drifting gratings during alternating 10 s blocks (Figure 3A). For the data shown in Figure 3 & Supplemental Figure 1, stimulus diameter alternated between 1° & 2°, or 1° & 12° in separate 5 min runs. For those in Supplemental Figure 3, Supplemental Figure 4, & Supplemental Figure 5, diameter alternated between 2° & 12°. Stimulus duration was 400 ms, with a 225 ms inter-stimulus interval. Sixteen gratings were presented at the center of the screen during each block; to prevent adaptation, gratings moved in one of 8 possible directions in a randomized and counter-balanced order. Twenty-five blocks (13 small, 12 large) were presented during each run. Stimulus contrast was either 3% or 98% (separate runs). Each subject completed 2-4 runs at each contrast level. Subjects performed a color / shape detection task at fixation during all fMRI experiments.

Functional localizer scans were acquired to facilitate ROI definition. Two different localizers were used; paradigm structure matched those above, except where noted. The first localizer was designed to identify human MT complex (hMT+); we use this notation to clarify that we did not attempt to distinguish areas MT & MST, both of which are motion selective^58^. Drifting and static gratings (15% contrast) were presented centrally in alternating blocks. Stimulus diameter was 1° (hMT+ data from Figure 3 & Supplemental Figure 1) or 2° (Supplemental Figure 3 & Supplemental Figure 4). The second localizer was used to identify regions of visual cortex that represented the smallest stimulus size^59^; checkerboard stimuli (100% contrast, phase-reversing at 8 Hz) were presented in center and surrounding regions in an alternating order across 16 blocks. Center diameter was either 1° (with a 1° gap; Figure 3 & Supplemental Figure 1) or 2° (Supplemental Figure 3, Supplemental Figure 4, & Supplemental Figure 5); inner and outer diameter of the surround annulus were always 2° and 12°, respectively.

FMRI data were processed in BrainVoyager (Brain Innovation, Maastricht, The Netherlands), including motion and distortion correction, high-pass filtering (cutoff = 2 cycles/scan), and anatomical alignment. No spatial smoothing or normalization were performed; all analyses were within-subjects & ROI-based. ROIs were identified from the localizer data using correlational analyses, with an initial threshold of *p* < 0.05 (Bonferroni corrected). ROIs were defined for each hemisphere in 2 anatomical regions: motion-selective hMT+ in the lateral occipital lobe (Supplemental Figure 1C), and the region of EVC selective for the retinotopic position of the center stimulus (near the occipital pole; Supplemental Figure 1A). ROI position was verified by visualization on an inflated and flattened model of the cortical white matter surface. The top 20 most-significant voxels (in functional space) within each hemisphere were selected for analysis. In a few cases, there were not 20 functional voxels within a hemisphere ROI that met the statistical threshold above. For these subjects, the threshold was relaxed until 20 voxels from the surrounding region were included. ROIs in all subjects satisfied an uncorrected, one-tailed significance threshold of *p* < 0.002. Center-selective hMT+ sub-ROIs were defined based on a correlation analysis of their time course data during the center-vs.-surround localizer scan, with a further inclusion criterion of *p* < 0.05 (one-tailed). This allowed us to examine the fMRI response to increasing stimulus size within hMT+ voxels that showed some selectivity for the retinotopic position of the center stimulus, in addition to significant motion selectivity. Sub-ROIs in hMT+ could not be identified in 2 subjects from the first fMRI experiment, and 6 subjects from the MRS & fMRI experiment.

Average time courses were extracted from each sub-ROI for further analyses in MATLAB using BVQXTools. Time course data were split into epochs spanning 4 s before to 4 s after each block. For each type of block (e.g., small and large stimuli), response baseline was determined by averaging the signal across all such epochs between 0-4 s prior to block onset. The time course in each epoch was then converted to percent signal change by subtracting and then dividing by the baseline, and multiplying by 100. Converted time courses were then averaged across hemispheres in each run, and across runs in each subject. The average signal change from 8-12 s after the onset of the block (the response peak) served as the measure of the fMRI response.

#### MR spectroscopy

Spectroscopy data were acquired using a MEGA-PRESS^60^ sequence (3 cm isotropic voxel, 320 averages of 2048 data points, 2 kHz spectral width, 1.4 kHz bandwidth refocusing pulse, VAPOR water suppression, 2 s TR, 68 ms TE). Editing pulses (14 ms) were applied at 1.9 ppm during “on” and 7.5 ppm during “off” acquisitions, interleaved every 2 TRs across a 16-step phase cycle. MRS data were collected within the following regions (Supplemental Figure 2): hMT+ in lateral occipital cortex, EVC in mid-occipital cortex, and a region of the central sulcus in parietal cortex known as the “hand knob”^61^. Voxels were positioned based on anatomical landmarks using a T_1_-weighted anatomical scan collected in the same session, while avoiding contamination by CSF, bone, and fat. The EVC voxel was placed medially within occipital cortex adjacent to the occipital pole, and aligned parallel to the cerebellar tentorium. The parietal voxel was centered on the “hand knob” within the central sulcus, and aligned parallel to the dorsolateral cortical surface. The hMT+ voxel was placed in the ventrolateral occipital lobe, parallel to the lateral cortical surface. Further positioning information for hMT+ was obtained using an abbreviated version of the hMT+ fMRI localizer described above (65 TRs, each 3 s, resolution 3x3x5 mm, 14 slices with 0.5 mm gap). These localizer data were processed on-line at the scanner, using a GLM analysis in the Philips iViewBOLD software to identify hMT+ voxels in lateral occipital cortex that responded more strongly to moving vs. static gratings (threshold *t* > 3.0). The hMT+ MRS voxels were centered on these functionally identified regions in the left and right hemispheres within each subject. To mitigate the detrimental effects of gradient heating during fMRI on the MRS data quality, the functional localizer data were acquired prior to the T_1_ anatomical scan. MRS data were acquired in both left & right hMT+ for all subjects. Two measurements in EVC were obtained for all subjects except 1. Both values were averaged for hMT+ and EVC. Parietal cortex was measured only once, and was not obtained in 1 subject.

To quantify the concentration of GABA+ within each voxel, MRS data were processed using the Gannet 2.0 toolbox^62^. We refer to this measurement as GABA+ to note that it includes some contribution from macromolecules that is not accounted for in our analysis^63^. Processing included automatic frequency and phase correction, artifact rejection (frequency correction parameters > 3 SD above mean), and exponential line broadening (3 Hz). The GABA+ peak was fit with a Gaussian (Supplemental Figure 2D & E), and the integral of the fit served as the concentration measurement. This GABA+ value was scaled by the integral of the unsuppressed water peak, fit with a mixed Gaussian-Lorentzian. The GABA+ value was corrected based on the concentration of gray and white matter using Equation 2 from^64^, assuming a ratio of GABA+ in white to gray matter (*α*) of 0.5. Gray and white matter concentrations were obtained for each voxel in each subject by segmenting the T_1_ anatomical data using SPM8^65^. This correction did not qualitatively affect our results. All MRS scans and corresponding psychophysical & fMRI data were collected within a maximum 2-week time-period, as previous work has shown GABA measurements are relatively stable across several days^24, 66^.

### Statistics

All statistical analyses were performed in MATLAB. *F*-test statistics were obtained using repeated measures ANOVAs (e.g., 4 repeated psychophysical threshold estimates per subject), with subjects treated as a random effect. Stimulus size was treated as a continuous variable where appropriate. To examine whether our data were normally distributed (as assumed by parametric tests such as an ANOVA), we manually inspected the distributions of our data in each condition, and used the Shapiro-Wilk test of normality. Cases in which deviations from normality were observed were further tested using the non-parametric equivalent of an ANOVA (Friedman’s test). The *p*-values obtained in all such cases were smaller for the Friedman’s test than for the corresponding ANOVA, thus we report the larger values and the more conventional statistic. Correlation values (*r*) are Pearson’s correlation coefficients (two-tailed, unless otherwise noted). Significant correlations were confirmed using a (non-parametric) permutation test, which involved randomly shuffling the data being correlated across subjects in each of 10,000 iterations. The proportion of permuted correlations whose absolute value was greater than that of the real correlation served as the measure of significance. Permutation tests consistently yielded smaller *p*-values than the corresponding Pearson’s correlations; the larger values are reported. Power analyses were performed to ensure that sample sizes were large enough such that the probability of type II error was less than 20%. Importantly, the sample sizes in our correlational analyses (n = 21 to 27) permit us to detect correlations of *r* ≥ 0.58 with the same power (probability of type II error less than 20%)^67^, assuming a two-tailed significance threshold of *p* = 0.05.

### Data and code availability

Data and custom analysis code will be made available by the authors upon request.

## Acknowledgements

We thank Geoffrey M. Boynton for providing the MATLAB functions for the model, and for comments on the manuscript. We also thank Brenna Boyd, Judy Han, Heena Panjwani, Micah Pepper, Meaghan Thompson, Anne Wolken, the UW Diagnostic Imaging Center, and the UW Investigational Drug Service for their help with subject recruitment and/or data collection. This work was supported by funding from the National Institute of Health (F32 EY025121 to MPS, R01 MH106520 to SOM, T32 EY007031, P41 EB015909 and R01 EB016089).

## Supplemental Information

In the Results and Methods, we present a summary of the standard normalization model (Equation 1), which is a direct application of the work from^19^. A more complete description is provided below. The parameters *E* and *S* represent the excitatory and suppressive drive within the model, which are a function of the spatial extent (*x*), orientation (*θ*), and contrast (*c*) of the stimulus, as well as the width of model tuning in space and orientation for both excitation (*x*_*w_e*_ , *θ*_*w_e*_) and suppression (*x*_*w_s*_ , *θ*_*w_s*_). This relationship can be expressed as:

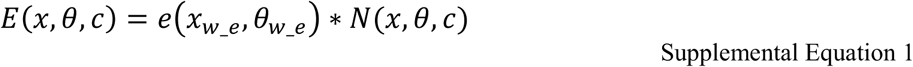

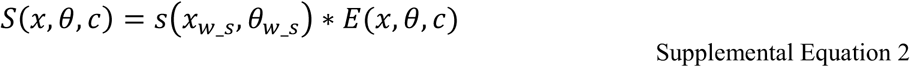

where *N* is a “neural image” that represents the population response to a given stimulus as a 2-dimensional Gaussian (*x* & *θ*), whose amplitude is set by *c*, and *\** denotes convolution. The terms *e* and *s* are also 2-D Gaussians that represent the selectivity (tuning width) of excitation and suppression, respectively. To predict the model response rate (*R*), the excitatory drive (*E*) is divided by the sum of the suppressive drive (*S*) and the semi-saturation constant (*σ*), as in Equation 1, which is reprinted here:

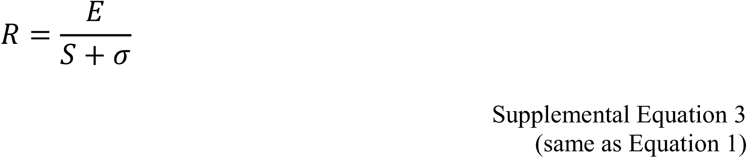

The predicted threshold (*T*) is determined according to the following procedure, which is summarized in Equation 3:

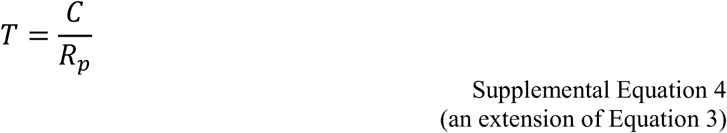

Where

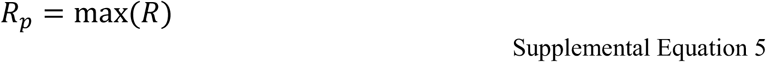

Thus, the threshold duration for motion discrimination predicted by the model (*T*) is a function of the peak response rate (*R*_*p*_) and the criterion response level (*C*) required for a perceptual judgment. We use a winner-take-all rule (Supplemental Equation 5) in determining *T*, which is consistent with studies in macaques showing that behavioral performance during motion direction discrimination may be accounted for by the response of a small pool of neurons whose tuning properties are well-matched to the stimulus (i.e., centered in space and orientation)^3, 56^.

Full parameters for each application of the model are given below in Supplemental Table 1. In the version of the model shown in Figure 4 that characterizes the results from the experiment using lorazepam, we include an additional parameter *A* that scales the response *R* from Supplemental Equation 3.

**Supplemental Table 1.** Normalization model parameters. Arbitrary units abbreviated as a.u. Lorazepam abbreviated LZ. Changes in contrast gain were modeled by varying stimulus contrast (e.g., stimulus contrast for the LZ model is the square root of the contrast for the placebo model). Response gain changes were modeled through the inclusion of a parameter that scaled the predicted response (*A*). For the MRS model, the effect of hMT+ GABA was modeled by varying the response criteria (lower value is 80% of the higher criterion).

| <u>Parameter name</u> | <u>Parameter value</u> |  |  |
| --- | --- | --- | --- |
|  | Basic model | Lorazepam model | hMT+ GABA<br>MRS model |
| Stimulus contrast | 0.03 or 0.98 | 0.03 or 0.98 (placebo),<br>0.017 or 0.099 (LZ) | 0.03 or 0.98 |
| Stimulus spatial center ($x$, a.u.) | 0 | 0 | 0 |
| Stimulus spatial width (a.u.) | 1, 2, or 12 | 0.84, 1.7, or 10 | 1, 2, or 12 |
| Stimulus orientation ($\theta$, °) | 90 | 90 | 90 |
| Stimulus orientation width (°) | 5 | 5 | 5 |
| Excitatory spatial<br>pooling width ($x_{w_e}$, a.u.) | 5 | 6.5 | 4.5 |
| Excitatory orientation<br>pooling width ($\theta_{w_e}$, °) | 25 | 5 | 25 |
| Suppressive spatial<br>pooling width ($x_{w_s}$, a.u.) | 40 | 40 | 15 |
| Suppressive orientation<br>pooling width ($\theta_{w_s}$, °) | 50 | 25 | 50 |
| Semi-saturation constant ($\sigma$, a.u.) | 0.0002 | 0.0001 | 0.0001 |
| Criterion (at 3% contrast, a.u.) | 300 | 775 | 300 (higher),<br>240 (lower) |
| Criterion (at 98% contrast, a.u.) | 650 | 775 | 375 (higher),<br>300 (lower) |
| Response scalar ($A$, a.u.) | N/A | 1 (placebo), 0.85 (LZ) | N/A |

**Supplemental Figure 1.**
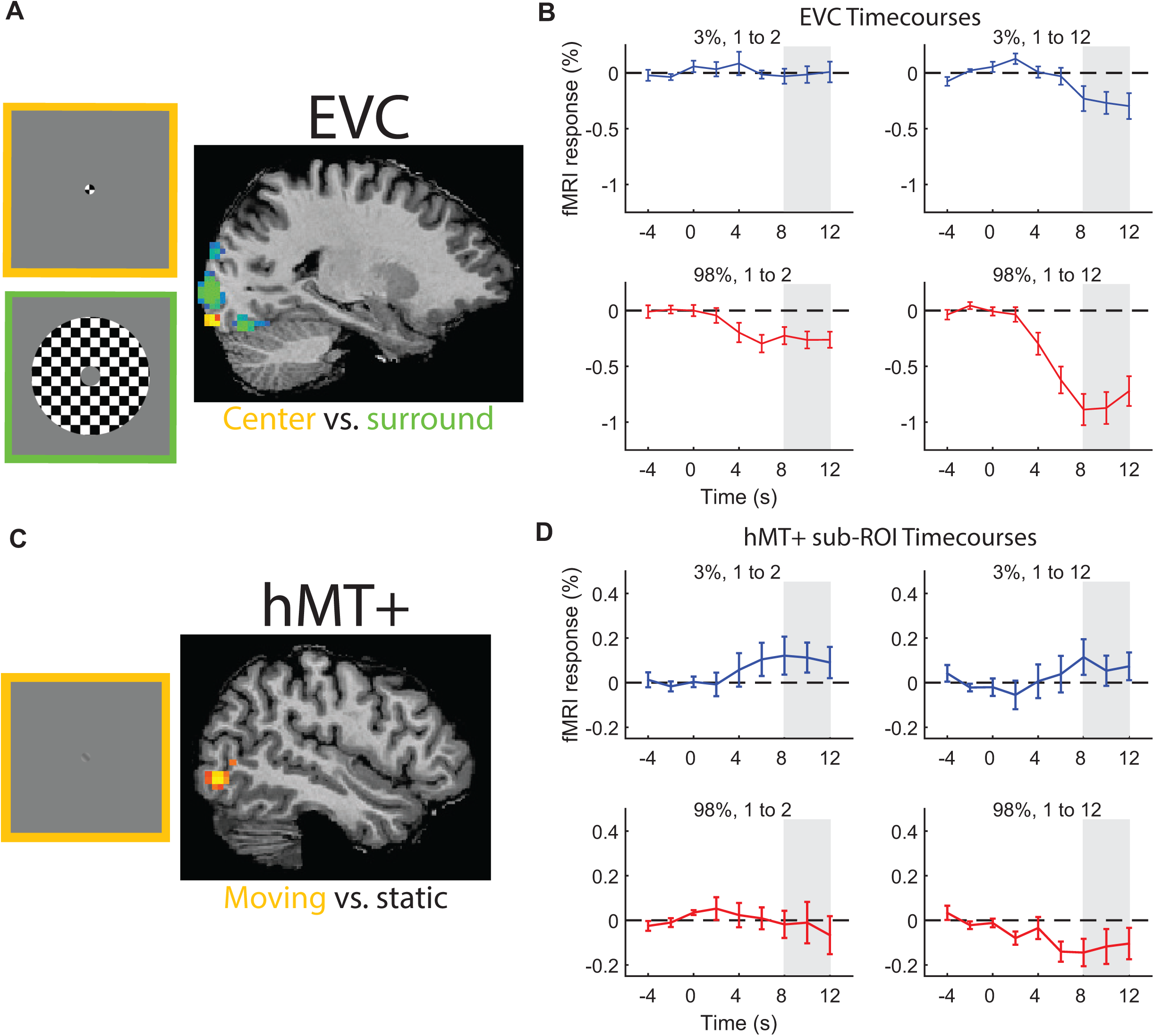
Regions-of-interest (ROIs) and response time courses for size-dependent fMRI responses. In **A**, center (orange) and surrounding (green) localizer stimuli are shown on the left. On the right, an example ROI in EVC was defined from voxels whose time course correlated positively with presentation of the center stimulus (orange; see Methods). Average time courses (**B**) from EVC ROIs for low (blue) and high contrast (red) stimuli from N = 10 subjects. At time = 0, stimulus size increased from 1 to 2° (left) or 1 to 12° (right). Gray region shows the time period where the peak response was calculated (shown in Figure 3). Error bars are mean ± s.e.m. An example of the hMT+ localizer stimulus (left) and the motion-selective ROI (right) are shown in **C**. Within this hMT+ ROI, a sub-region was identified that also responded selectively to the center > surround localizer (from **A**, see Methods). Sub-ROIs were identified in N = 8 subjects. Response time courses for hMT+ sub-ROIs (**D**) are shown, as in **B**.

**Supplemental Figure 2.**
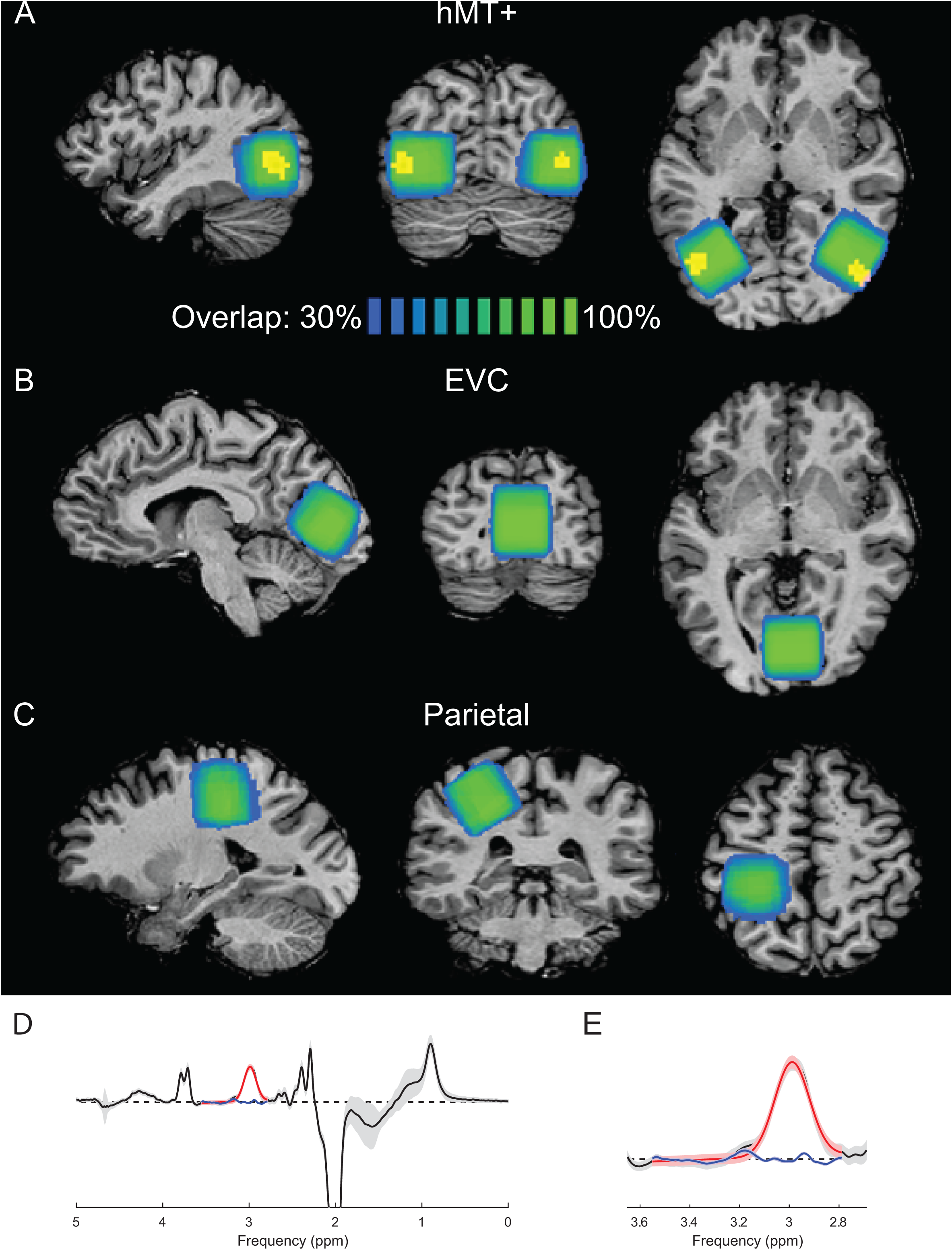
MRS voxel placement and GABA+ fitting. Mean spectroscopy voxel locations in right and left hMT+ (**A**) are shown in blue – green (color indicates % overlap across subjects). To visualize positioning across individuals, each subject’s voxel coordinates were mapped to Talairach space, percent overlap was calculated, and the result was projected onto a representative anatomical image. Note that this is only for visualization purposes; the exact voxel placement for each subject was dictated by individual anatomy. Average location of hMT+ across subjects is shown in yellow (determined using a similar method as average MRS voxel position, correlation threshold > 0.3). Images are shown in neurological convention (i.e., left is left). Voxel location for EVC (**B**) and Parietal cortex (**C**) are also shown. (**D**) Average spectrum across subjects from left hMT+ is shown in black (error bars show SD). Average fit to the GABA peak (red) and residuals (blue) are also shown. (**E**) Zoomed region from **D**.

**Supplemental Figure 3.**
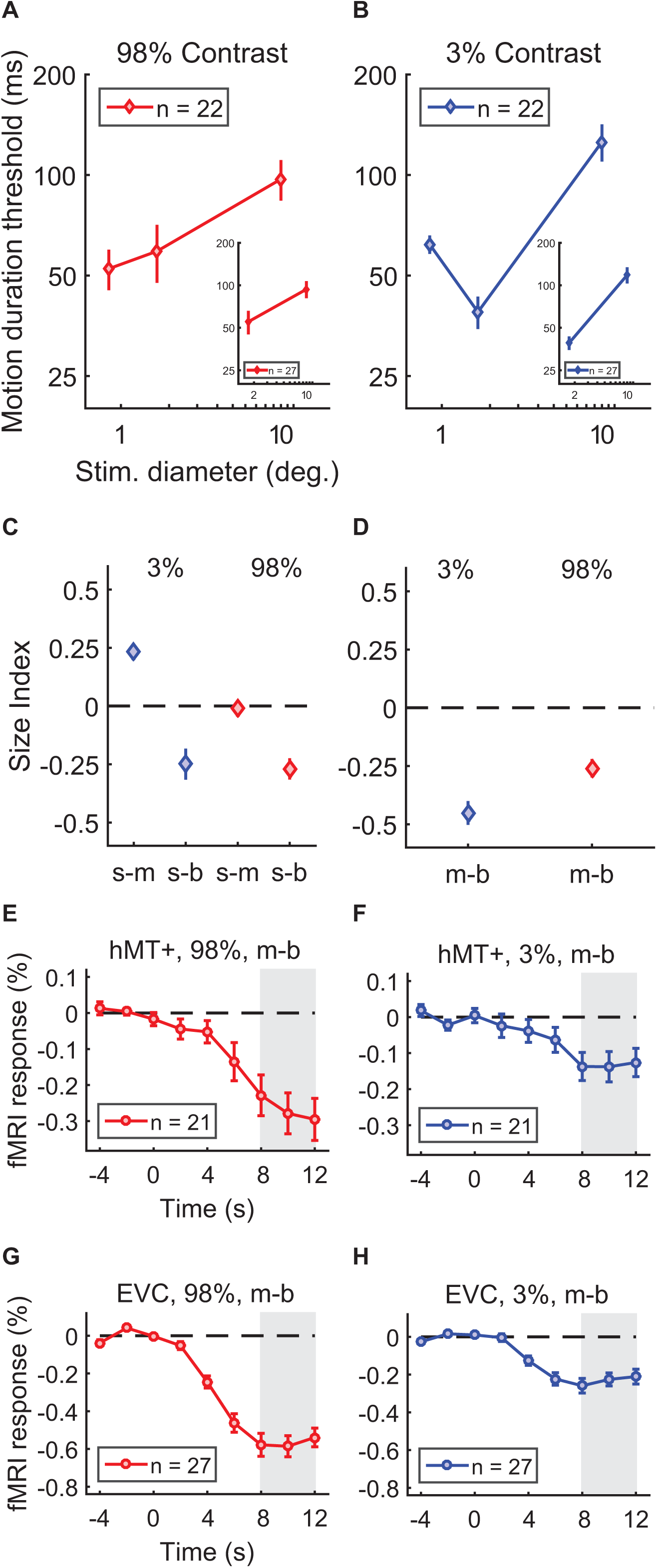
Motion discrimination thresholds are shown for the subjects who participated in all 6 stimulus conditions in **A** & **B** (N = 22). Thresholds were obtained from 5 additional subject for only the medium and big stimulus sizes; inset panels show the data from all subjects (N = 27) in these 4 conditions. Size indices are shown for small-medium and small-big comparisons in **C** (N = 22), and medium-big in **D** (N = 27). All subjects (N = 27) participated in an fMRI experiment that measured the response to increasing stimulus size. Response time courses for high (**E** & **G**) and low contrast (**F** & **H**) stimuli are plotted for regions representing the smallest stimulus size (**E** & **F**: hMT+ sub-ROIs, identified in N = 21 subjects; **G** & **H**: EVC ROIs, N = 27). Gray region shows the time period where the peak response was calculated (shown in Supplemental Figure 4G-H & Supplemental Figure 5E-F). Error bars are mean ± s.e.m.

**Supplemental Figure 4.**
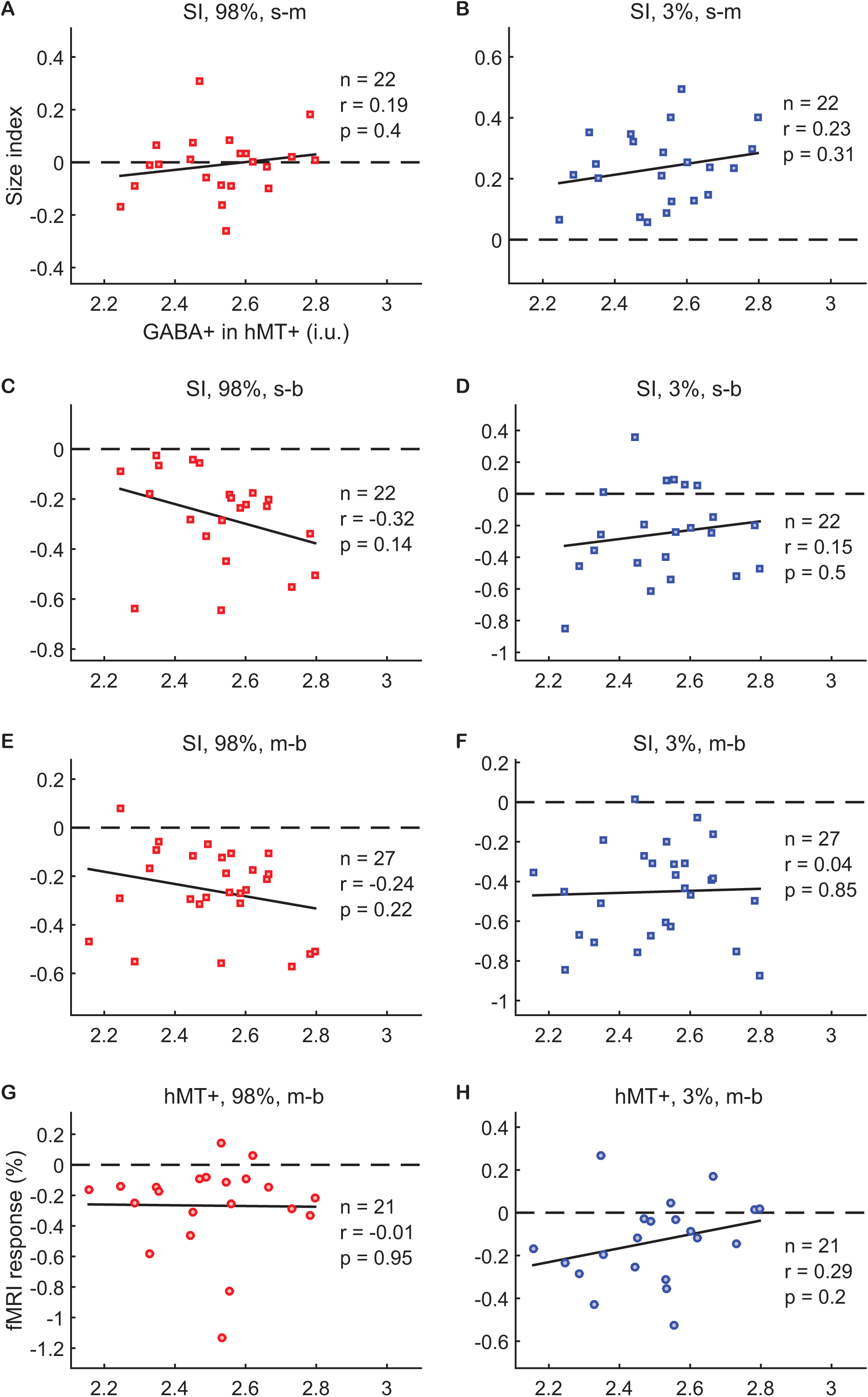
No relationship between GABA+ in hMT+ and suppression or summation. In all panels, the concentration of GABA+ in hMT+ (left and right measured in separate scans and averaged) is shown for each subject along the x-axis. Psychophysical size indices are shown along the y-axis in panels **A**-**F**, which quantify the change in motion discrimination thresholds for small-medium (s-m; **A** & **B**), small-big (s-b; **C** & **D**), and medium-big (m-b; **E** & **F**) comparisons. In **G** & **H**, the magnitude of the fMRI response in hMT+ evoked by increasing stimulus size (m-b) is shown along the y-axis. Stimulus contrast was 98% in panels **A**, **C**, **E**, & **G**, and 3% in **B**, **D**, **F**, & **H**. In all panels, negative values indicate suppression; positive values indicate facilitation. The number of subjects (n), correlation coefficient (r) and significance (p) is shown for each comparison. No significant correlations between hMT+ GABA+ and size indices or fMRI suppression in hMT+ were observed. These sample sizes are sufficient to detect correlations of *r* ≥ 0.58 (for n = 21) to *r* ≥ 0.52 (n = 27) with 80% power (probability of type II error < 20%)^67^.

**Supplemental Figure 5.**
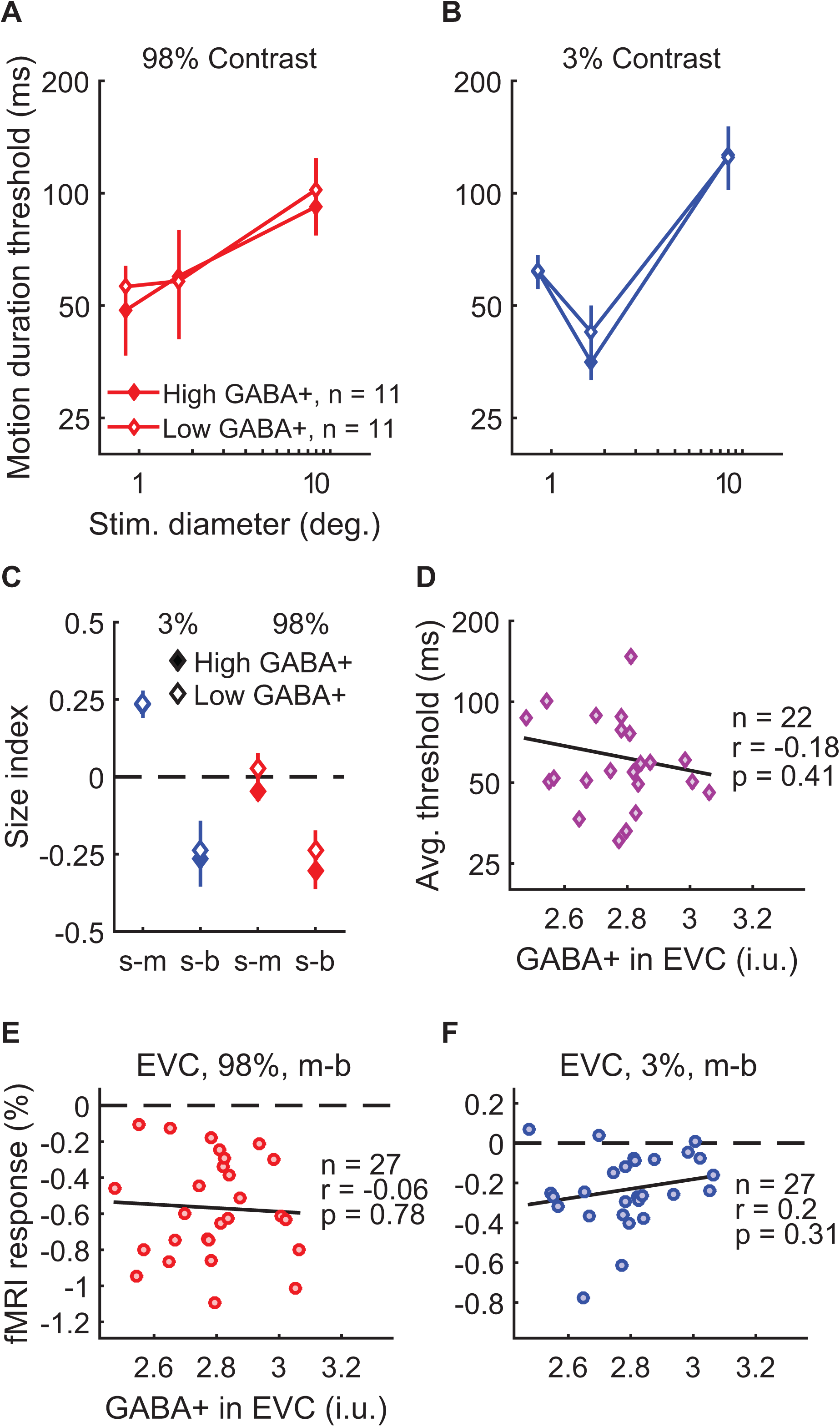
Examining task performance in terms of GABA+ in EVC. Thresholds (**A** & **B**) and size indices (**C**) are shown for subjects with lower (open symbols, N = 11) and higher GABA+ (filled symbols, N = 11; groups defined by median split) in EVC (average of 2 measurements). Error bars are mean ± s.e.m. GABA+ in EVC had no significant relationship with psychophysical size indices (correlation with 3% s-m: *r*_20_ = 0.11, *p* = 0.61; 3% s-b: *r*_20_ = -0.05, *p* = 0.84; 3% m-b [not shown]: *r*_25_ = 0.01, *p* = 0.96; 98% s-m: *r*_20_ = -0.04, *p* = 0.87; 98% s-b: *r*_20_ = -0.23, *p* = 0.30; 98% m-b [not shown]: *r*_25_ = -0.17, *p* = 0.39). As shown in **D**, there was also no relationship between GABA+ in EVC and overall motion discrimination performance (average thresholds are the geometric mean of all 6 stimulus conditions). In **E** & **F**, fMRI responses in EVC to increasing stimulus size (medium-big) are shown along the y-axis. No significant correlations with GABA+ in EVC were observed. These sample sizes are sufficient to detect correlations of *r* ≥ 0.57 (for n = 22) to *r* ≥ 0.52 (n = 27) with 80% power (probability of type II error < 20%)^67^.

**Supplemental Figure 6.**
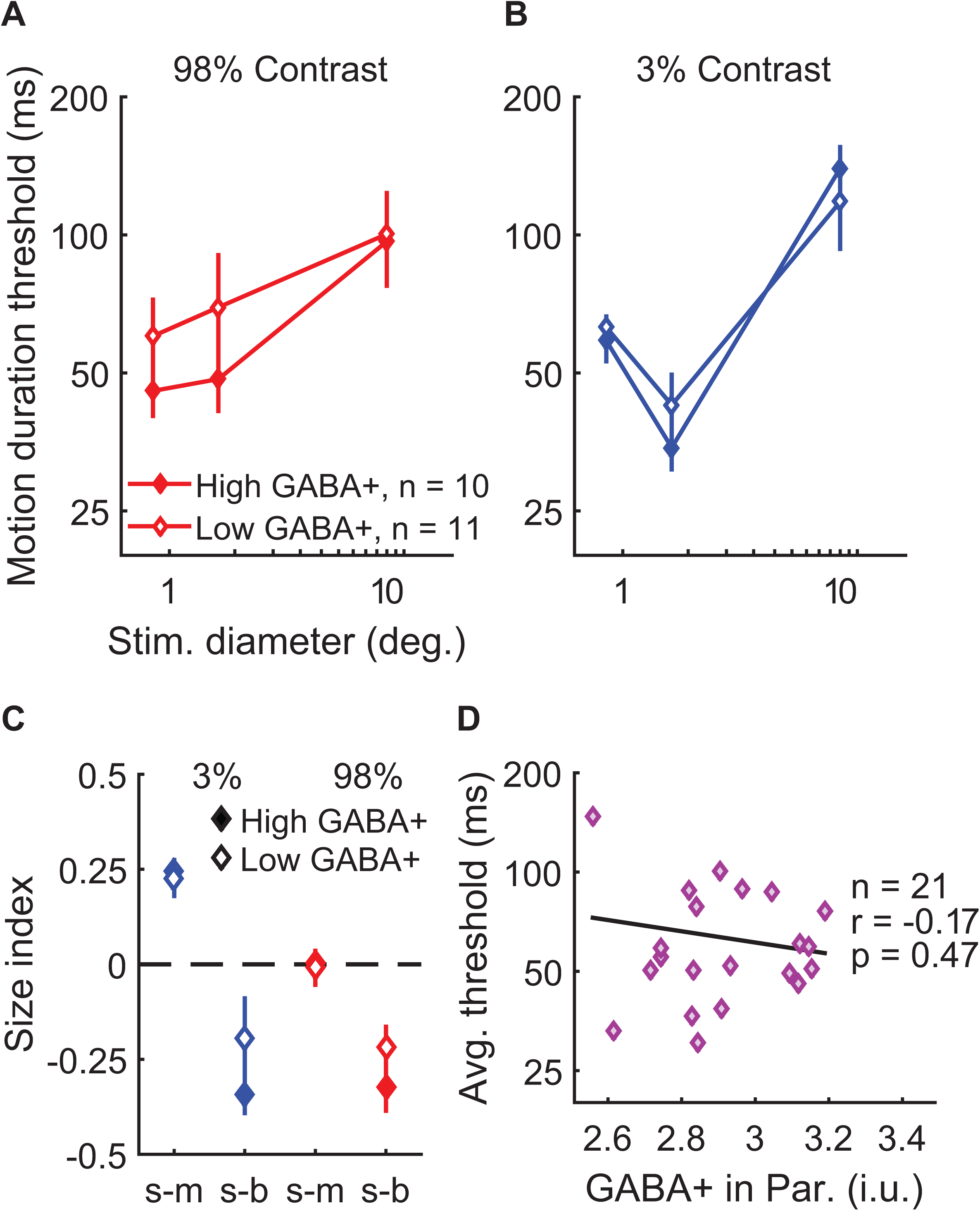
Examining task performance in terms of GABA+ in parietal cortex. Thresholds (**A** & **B**) and size indices (**C**) are shown for subjects with lower (open symbols, N = 11) and higher GABA+ (filled symbols, N = 10; groups defined by median split) in parietal cortex (Par.). Error bars are mean ± s.e.m. There was no relationship between Par. GABA+ and psychophysical size indices (correlation with 3% s-m: *r*_19_ = 0.27, *p* = 0.23; 3% s-b: *r*_19_ = -0.10, *p* = 0.66; 3% m-b [not shown]: *r*_24_ = -0.15, *p* = 0.46; 98% s-m: *r*_19_ = 0.10, *p* = 0.67; 98% s-b: *r*_19_ = -0.28, *p* = 0.21; 98% m-b [not shown]: *r*_25_ = -0.16, *p* = 0.44). Nor was there a significant relationship between Par. GABA+ and overall motion discrimination performance (**D**; average thresholds are the geometric mean of all 6 stimulus conditions). These sample sizes are sufficient to detect correlations of *r* ≥ 0.57 (for n = 21) to *r* ≥ 0.53 (n = 26) with 80% power (probability of type II error < 20%)^67^.

**Supplemental Figure 7.**
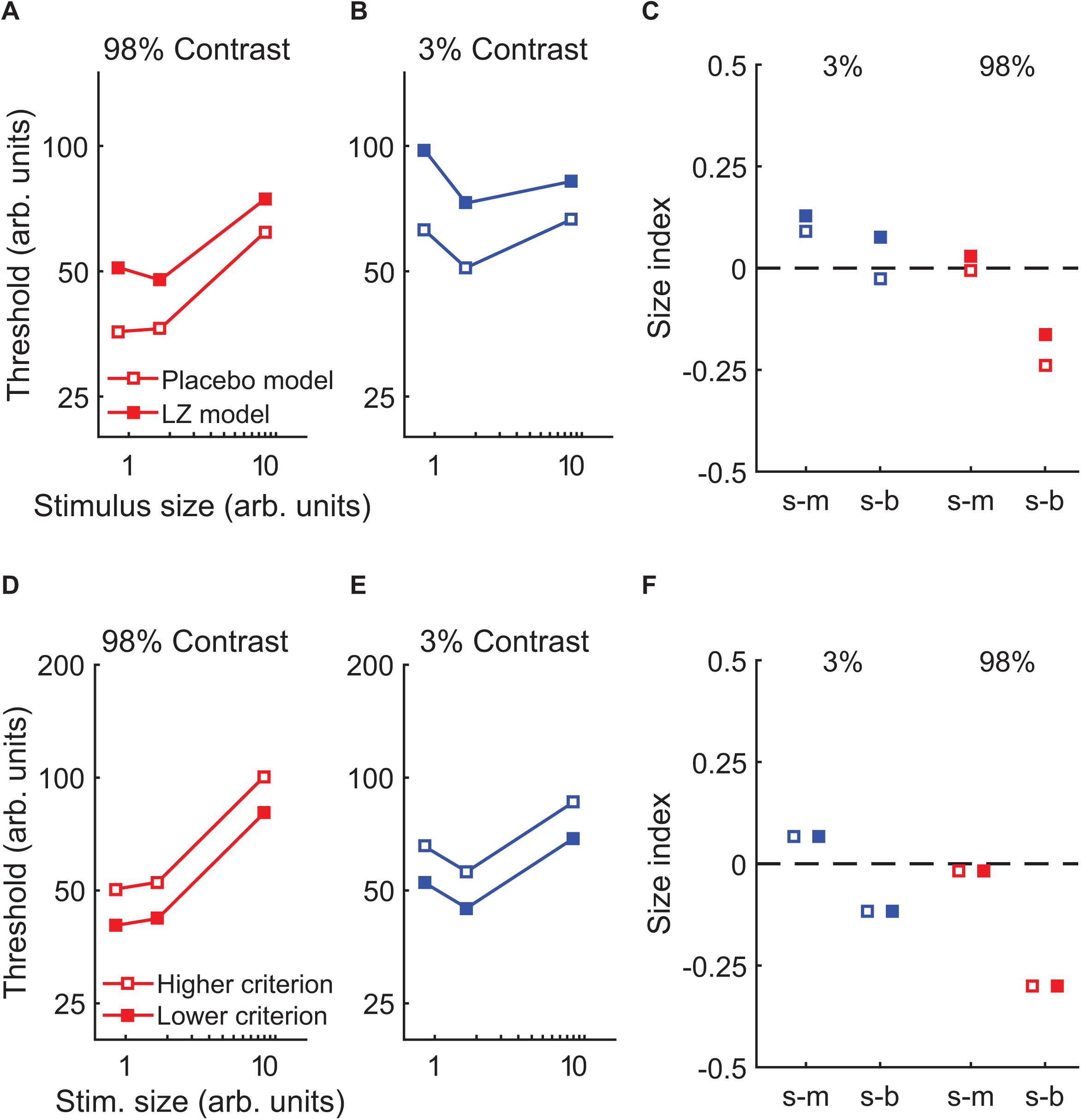
Describing the effects of GABA during motion discrimination using the normalization model. The effect of lorazepam (LZ) versus placebo was described by reducing both contrast and response gain within the normalization model (Supplemental Table 1). Using this model, we were able to predict the effect of LZ on motion discrimination thresholds (**A** & **B**) and size indices (**C**), in a manner which mirrors the data from Figure 4. The relationship between GABA+ in hMT+ (measured using MRS) and motion discrimination performance was modeled by a change in behavioral response criteria (Supplemental Table 1). Lowering the criterion leads to lower predicted thresholds overall (**D** & **E**), but does not affect size indices (**F**). The predictions of this model are well matched to the data shown in Figure 5. In panel **F**, indices for higher & lower criterion models are offset along the x-axis, to prevent overlap and illustrate that the values are identical.

